# Genome-guided, field-deployable loop-mediated isothermal amplification (LAMP) assay for specific detection of *Dickeya dadantii*

**DOI:** 10.1101/2024.05.04.592507

**Authors:** Stefania Montesinos, Garima Tyagi, Zhuokun Feng, Ella Hampson, Achyut Adhikari, Michael Minaai, Landon Wong, Meagan Haubner, Shefali Dobhal, Dario Arizala, Sharon A. Andreason, Dimitre Mollov, Francisco Ochoa-Corona, Jon-Paul Bingham, Jenee Odani, Daniel Jenkins, Li Maria Ma, Jacqueline Fletcher, James P. Stack, Mohammad Arif

**Affiliations:** Department of Plant and Environmental Protection Sciences, University of Hawaii at Manoa, Honolulu, HI, USA; Department of Molecular Biosciences and Bioengineering, University of Hawaii at Manoa, Honolulu, HI, USA; Department of Tropical Plant and Soil Sciences, University of Hawaii at Manoa, Honolulu, HI, USA; USDA Agricultural Research Service, U.S. Vegetable Laboratory, Charleston, SC; USDA Agricultural Research Service, Horticultural Crops Disease and Pest Management Research Unit, Corvallis, OR; Institute for Biosecurity & Microbial Forensics, Oklahoma State University, Stillwater, OK, USA; Department of Human Nutrition, Food and Animal Sciences, University of Hawaii at Manoa, Honolulu, HI, USA; Department of Plant Pathology, Kansas State University, Manhattan, KS, USA

**Author notes:** These authors contributed equally to this work (Stefania Montesinos, Garima Tyagi, Zhuokun Feng, Ella Hampson, Achyut Adhikari, Michael Minaai, Landon Wong, Meagan Haubner).

**Keywords:** *Dickeya dadantii*, LAMP, soft rot, phytobacteria, diagnostics

## Abstract

Potatoes, among the most economically significant crops worldwide, are susceptible to various plant pathogens that significantly impact their propagation, production, storage, and distribution. Soft rot disease, caused primarily by *Dickeya* and *Pectobacterium*, results in substantial economic losses to the agricultural industry annually. In this study, we developed a rapid, reliable, and field-deployable loop-mediated isothermal amplification (LAMP) assay for detecting *D. dadantii*, a common soft rot causing bacteria. The *D. dadantii*-specific LAMP primers were designed targeting a highly conserved genomic region within *D. dadantii*, the TetR/AcrR family transcriptional regulator CDS and its flanking sequences. This assay was thoroughly validated with the members of inclusivity (nine strains of *D. dadantii*) and exclusivity panels (85 strains, including all *Dickeya* species, related taxa, and host DNA), detecting no false positives or negatives. The limit of detection (LOD) was established by performing assays with 10-fold serially diluted pure gDNA of *D. dadantii* and gDNA spiked with host crude extract; the assay detected the target pathogen down to 1 pg (188 copies) without being adversely affected by the host crude extract. The developed LAMP assay specifically detected the target pathogen in infected plant materials. Additional multi-operator blind and multi-instrument tests were conducted to assess the assay’s robustness and applicability, consistently yielding accurate results without false positives or negatives. These findings demonstrate the assay’s potential utility for biosecurity, routine diagnostics, and epidemiological studies.

## INTRODUCTION

As a consequence of the SARS-CoV-2 (COVID-19) viral pandemic, 83 to 132 million people were unable to secure food necessary to meet their nutritional needs in 2020 (HLPE 2020). Historically, the potato (*Solanum tuberosum* L.) has played a significant role in alleviating food insecurity during times of crisis (Campos and Ortiz, 2020). Its adaptability, high yield potential, and nutritional value make the potato the world’s most important dicot food crop for human consumption (CIP 2022). Potatoes rank as the fifth most economically significant crop, with a global production of approximately 380 million tons in 2016 (Youdkes et al., 2020). A substantial loss of potato yields due to disease could lead to a considerable economic downturn and exacerbate the current food crisis (Savary et al. 2012).

Potato soft rot and blackleg are among the most prevalent diseases affecting potatoes (Arif et al., 2022; Boluk et al, 2019; Cardoza et al, 2016; Czajkowski et al. 2012). The genus *Dickeya*, within the Pectobacteriaceae family, includes 12 species and one subspecies of bacterial plant pathogens (Arif et al, 2022; Hugouvieux-Cotte-Pattat et al, 2021; Wang et al, 2020). These pathogens are capable of causing soft rot diseases in a wide range of monocotyledonous and dicotyledonous crops, with potato tubers being one of the most commonly affected (Arif et al., 2022; Boluk et al. 2021; He et al. 2021; Liu et al. 2016; Soleimani-Delfan, 2015). Due to its broad host range and rapid infection rate, *Dickeya* ranks as one of the most detrimental bacterial plant pathogens (Mansfield et al. 2012). *Dickeya* species infect hosts through natural openings or wounds in the plant tissue and produce plant cell wall degrading enzymes (PCWDEs), which macerate the plant’s cellular structure, leading to soft rot (DeLude et al. 2022; Helmann et al. 2022). *Dickeya dadantii* is recognized as the ninth most economically significant bacterial plant pathogen worldwide (Mansfield et al. 2012). Unfortunately, the diagnostic methods available for identifying these pathogens are currently limited (He et al. 2021; Pritchard et al, 2013). The agricultural destruction caused by bacterial infections in potato crops underscores the critical need for developing accurate, sensitive, field-deployable diagnostic assays to detect plant pathogens and effectively control their spread (Dobhal et al, 2024; DeLude et al. 2022; Boluk et al, 2020).

While diagnostic assays have been developed for *Dickeya* species, there is currently a lack of comprehensive, cost-effective, field-deployable assays for *D. dadantii* (Dobhal et al. 2020, 2024; Boluk et al. 2020; Ocenar et al. 2019; Safenkova et al. 2017). The molecular technologies used to detect *D. dadantii* currently rely on laboratory settings that require expensive equipment and highly skilled personnel, presenting significant challenges for point-of-care diagnosis (He et al. 2021). Polymerase chain reaction (PCR) and quantitative real-time PCR (qPCR) methods often exhibit limited accuracy and sensitivity due to their low tolerance to host inhibitors, resulting in false negatives (Arif et al. 2021). Isothermal methodologies, such as loop-mediated isothermal amplification (LAMP) and recombinase polymerase amplification (RPA), on the other hand, offer efficient detection capabilities, enabling the identification of pathogens with standard laboratory equipment within 30 minutes (Dobhal et al, 2024; Arif et al. 2021; Ivanov et al. 2021; Boluk et al. 2020). LAMP, in particular, stands out as the most cost-effective option for pathogen detection, capable of amplifying target DNA up to 10^9^ copies using a simple water bath (Notomi et al. 2000; Zou et al. 2020). By utilizing a combination of four to six primers and DNA polymerase, it enables the simultaneous amplification of target DNA into millions of copies of stem-loop structured products (Notomi et al. 2000; Parida et al. 2008). This amplified DNA can be visualized through various mechanisms, including visual turbidity, fluorescence, agarose gel electrophoresis, and real-time monitoring (Lin et al. 2022; Domingo et al. 2021). The user-friendly application and versatility of LAMP assays make them an ideal diagnostic tool for both high and low-resource settings in the detection and diagnosis of *D. dadantii* (Parida et al. 2008). Identifying a unique genomic target for primer design can make the developed assay robust and enhance its adaptability across various laboratories and field settings (Arif et al., 2021; Domingo et al., 2021; DeLude et al., 2022).

This research aimed to develop a LAMP assay for the specific detection of *D. dadantii*. By conducting comparative genomic analyses between *D. dadantii* and closely related species, we identified a unique genomic region exclusive to *D. dadantii*. From this region, we designed LAMP primers to enable rapid identification and diagnosis of *D. dadantii* strains. This diagnostic assay offers significant advantages, including in-field pathogen detection, cost-effectiveness compared to traditional methods, and ensuring comprehensive coverage of diverse *D. dadantii* strains.

## RESULTS

### Target Region and *In-silico* Specificity

To develop a highly accurate and reliable assay, we identified a genome region exclusive to *D. dadantii*. This region includes the TetR/AcrR family transcriptional regulator CDS and its flanking sequence, spanning 121 base pairs. Primer sequences designed from this region showed no matches with any non-target species in the NCBI database, as detailed in Table 1. Testing these primers against the NCBI database confirmed their high specificity to all known strains of *D. dadantii*. The uniqueness of this selected region is illustrated in Figure 1.

**Figure 1.**
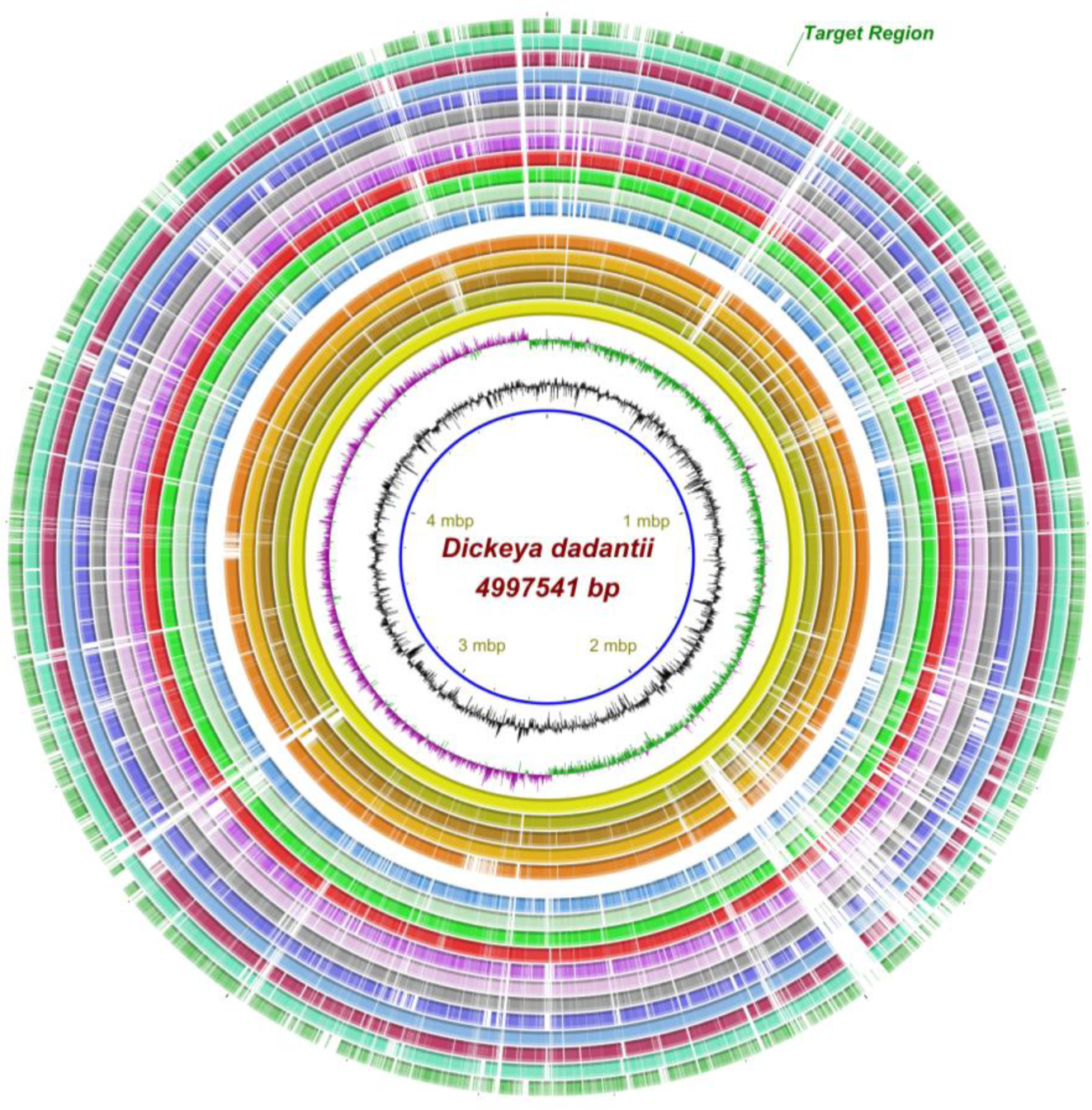
BLAST ring image generator (BRIG) plot illustrating the exclusive presence of the target region, TetR/AcrR family transcriptional regulator CDS, and its flanked sequences in the genome of *D. dadantii*. The ring image shows a multiple genome alignment comprising five *D. dadantii*, followed by eleven *Dickeya* species and one *Pectobacterium* species. The innermost ring represents the reference genome coordinates, *D. dadantii* DSM1820 (NZ_CP023467), in megabase pairs (Mbp). Black and alternating green/purple rings indicate the GC content ratio and GC skew (purple for positive and green for negative), respectively. The five inner lines, shaded from yellow to orange, correspond to BLASTn pairwise comparisons of *D. dadantii* strains DSM1820ᵀ (NZ_CP023467), 3937 (NC_014500), FZ06 (NZ_CP094943), M2-3 (NZ_CP077422), and S3-1 (NZ_CP076386), illustrating the conservation of the TetR/AcrR family transcriptional regulator CDS and its flanking region exclusively within these *D. dadantii* strains, highlighted in green. The outer 12 rings display genome alignments for *D. aquatica* 174/2ᵀ (NZ_LT615367), *D. chrysanthemi* Ech1591 (NC_012912), *D. dianthicola* ME23 (NZ_CP031560), *D. fangzhongdai* DSM101947ᵀ (NZ_CP025003), *D. lacustris* S29ᵀ (QNUT00000000), *D. oryzae* ZYY5ᵀ (SULL00000000), *D. parazeae* Ech586 (NC_013592), *D. poaceiphila* NCPPB 569ᵀ (NZ_CP042220), *D. solani* IPO 2222 (NZ_CP015137), *D. undicola* 2B12 (JSYG00000000), *D. zeae* PL65 (NZ_CP040817), and *P. brasiliense* 1692 (NZ_CP047495).

**Table 1.**
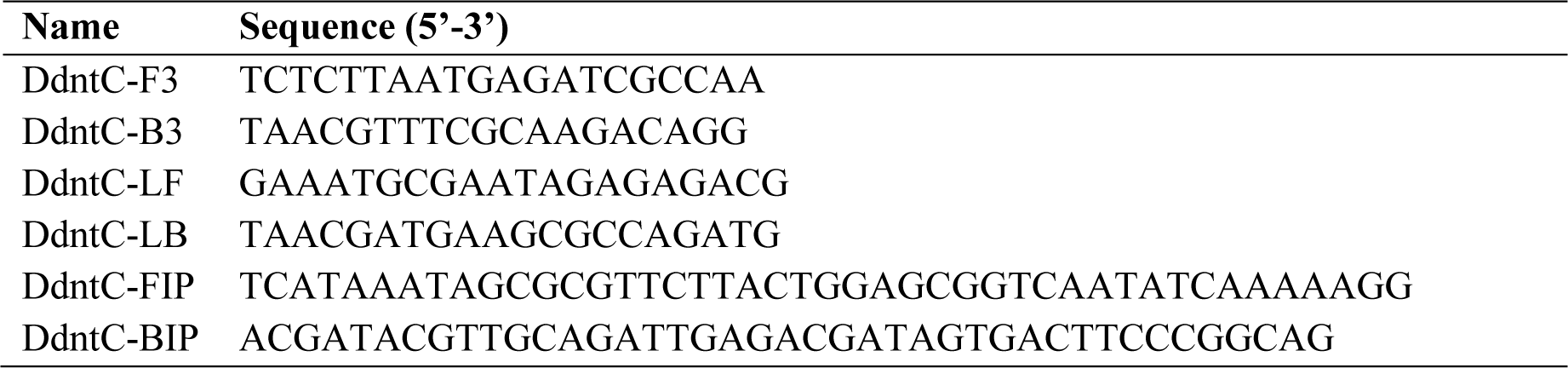
Sequences of loop-mediated isothermal amplification (LAMP) primers designed for specific and rapid detection of *Dickeya dadantii*.

### Specificity with Strains Included in Inclusivity and Exclusivity Panels

Strain inclusivity and exclusivity panels were utilized to rigorously validate the specificity of the LAMP assay. The inclusivity panel consisted of eight *D. dadantii* strains. The exclusivity panel was composed of closely related bacterial species, including *D. solani* (A5582), *D. oryzae* (A5417), *D. aquatica* (LMG27354), *D. fangzhongdai* (CFPB8607), *D. zeae* (A6066), *D. chrysanthemi* (A5415), and other species from closely related genera (Table 2). The LAMP assay tested negative with non-target bacteria and the DNA from plants or soil, in the exclusivity panel. However, amplification was observed with all eight strains of *D. dadantii* in the inclusivity panel, as shown in Figure 2. No false positives or negatives were detected at any stage of validation. All negative and positive results were confirmed in real-time PCR and visually through SYBR Green fluorescence.

**Figure 2.**
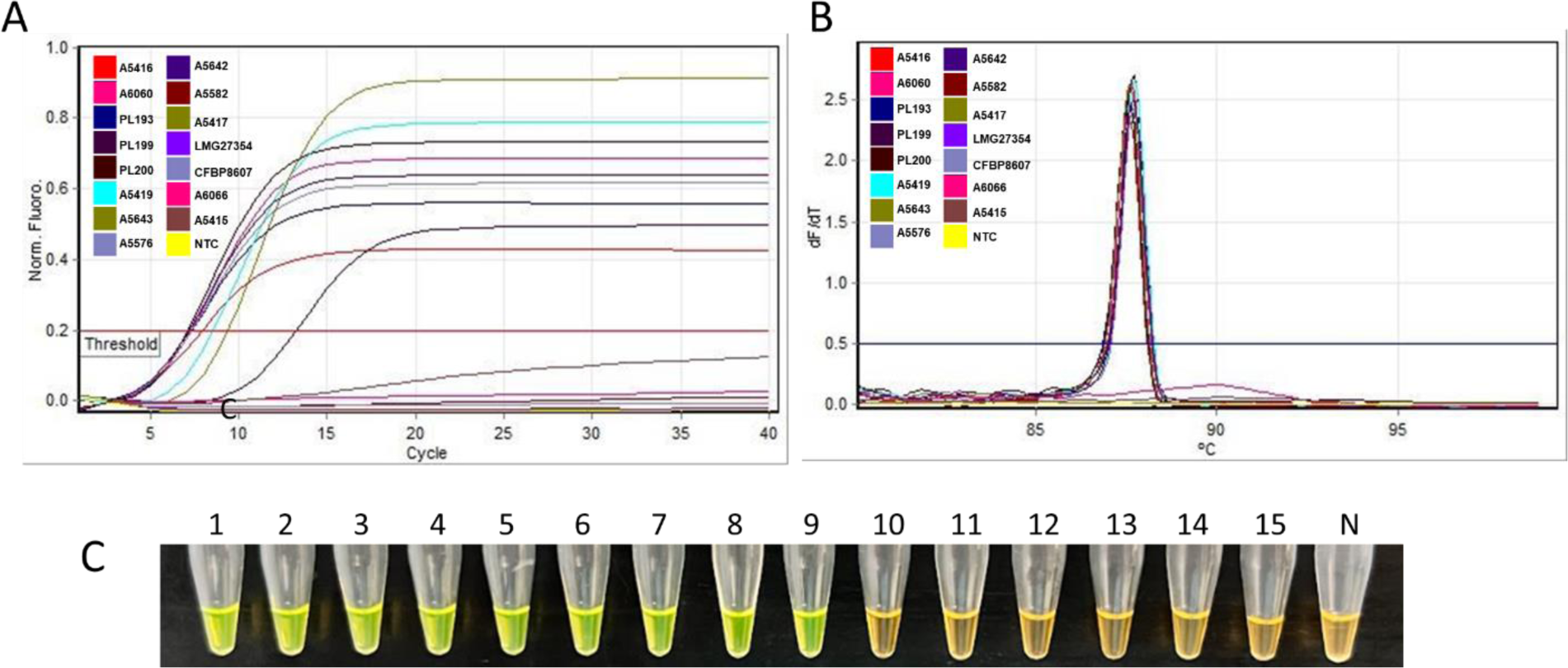
Specificity validation of the loop-mediated isothermal amplification (LAMP) assay designed for specific detection of *Dickeya dadantii*. (A) Sigmoidal amplification curves generated by nine *D. dadantii* strains comprising the inclusivity panel: CFBP1269T (positive control), CFBP3698, PL193, PL199, PL200, A5419, A5643, A5576, and A5642. No amplification occurred with genomic DNA (gDNA) from strains in the exclusivity panel or with the non-template control (NTC; water). (B) The melt curve analysis. (C) Colorimetric detection of LAMP reaction products under ambient light following the addition of SYBR Green I dye; positive and negative results are indicated by green and orange colors, respectively. Tube 1: CFBP8607^T^; tubes 2-9: *D. dadantii* strains CFBP3698, PL193, PL199, PL200, A5419, A5643, A5576, and A5642; tubes 10-15 contain representative strains from the exclusivity panel: A5582 (*D. solani*), A5417 (*D. oryzae*), LMG27354 (*D. aquatica*), CFPB8607 (*D. fangzhongdai*), A6066 (*D. zeae*), A5415 (*D. chrysanthemi*); and tube N: NTC (water).

**Table 2.** List of bacterial strains, plant hosts, and soil samples included in the inclusivity and exclusivity panels for the validation of *Dickeya dadantii* loop-mediated isothermal amplification (LAMP) assay. Hyphens (-) indicate data not available. “^T^” indicates a type strain.

| Species Name | Strain ID | Location | Host/Source | LAMP Results |
| --- | --- | --- | --- | --- |
| **Inclusivity Panel** |  |  |  |  |
| *Dickeya dadantii* | CFBP1269<sup>T</sup>; A5416 | Comoros, Africa | *Pelargonium capitatum* | Positive |
| *D. dadantii* ssp. *dadantii* | CFBP3698; A6060 | Cuba | *Musa* spp. | Positive |
| *D. dadantii* | PL193 | Hawaii, USA | *Solanum tuberosum* | Positive |
| *D. dadantii* | PL199 | Hawaii, USA | *S. tuberosum* | Positive |
| *D. dadantii* | PL200 | Hawaii, USA | *S. tuberosum* | Positive |
| *D. dadantii* ssp. *dieffenbachiae* | CFBP2051<sup>T</sup>; A5419 | USA | *Dieffenbachia* spp. | Positive |
| *D. dadantii* | CFBP6467; A5643 | Martinique, France | *Musa* spp. | Positive |
| *D. dadantii* | PRI2120; A5576 | Comoros, Africa | *P. capitatum* | Positive |
| *D. dadantii* | CFBP3855; A5642 | France | *Saintpaulia ionantha* | Positive |
| **Exclusivity Panel** |  |  |  |  |
| *Agrobacterium* sp. | A2961 | New York, USA | *Prunus avium* | Negative |
| *Clavibacter sepedonicus* | A2041 | Denmark | *S. tuberosum* | Negative |
| *C. phaseoli* | A6135 | Spain | *Phaseolis vulgaris* | Negative |
| *C. sepedonicus* | A6172 | Canada | *S. tuberosum* | Negative |
| *Candidatus Pectobacterium maceratum* | A1838 | - | *S. tuberosum* | Negative |
| *Curtobacterium flaccumfaciens pointsettiae* | A1147 | USA | *Euphorbia pulcherrima* | Negative |
| *C. flaccumfaciens* | A6267 | Hawaii, USA | Tulipa | Negative |
| *Dickeya aquatica* | LMG27354 | United Kingdom | River water | Negative |
| *D. chrysanthemi* | CFBP1270; A5641 | Denmark | *Parthenium argentatum* | Negative |
| *D. chrysanthemi* | CFBP2048; A5415 | USA | *Chrysanthemum morifolium* | Negative |
| *D. chrysanthemi* | A5468 | Hawaii, USA | Irrigation water | Negative |
| *D. chrysanthemi* | PL196 | Hawaii, USA | *S. tuberosum* | Negative |
| *D. chrysanthemi* | PL198 | Hawaii, USA | *S. tuberosum* | Negative |
| *D. dianthicola* | PL23 | Oahu, Hawaii | *S. tuberosum* | Negative |
| *D. dianthicola* | PL24 | Oahu, Hawaii | *S. tuberosum* | Negative |
| *D. dianthicola* | PL25 | Oahu, Hawaii | *S. tuberosum* | Negative |
| *D. dianthicola* | CFBP1200; A5418 | United Kingdom | *Dianthus caryophyllus* | Negative |
| *D. dianthicola* | PL195 | Hawaii, USA | *S. tuberosum* | Negative |
| *D. fangzhongdai* | CFBP8607 | China | Pear tree (bleeding cancer) | Negative |
| *D. lacustris* | LMG30899 | France | Water | Negative |
| *D. oryzae* | A5417 | Hawaii, USA | *Ananas comosus* | Negative |
| *D. solani* | PRI 2188; A5582 | Israel | *S. tuberosum* | Negative |
| *D. solani* | LMG25990 | Netherlands | *Hyacinthus orientalis* | Negative |
| *D. solani* | PRI 2187; A5581 | Israel | *S. tuberosum* | Negative |
| *D. solani* | PR1 2188; A5582 | Israel | *S. tuberosum* | Negative |
| *D. solani* | LMG25865; A6289 | Belgium | *S. tuberosum* | Negative |
| *D. undicola* | LMG30903 | Petaling Jaya, Malaysia | Fresh water | Negative |
| *D. zeae* | CFBP1889; A6066 | Malaysia | *A. comosus* | Negative |
| *D. zeae* | CFBP1890; A6067 | Malaysia | *A. comosus* | Negative |
| *D. zeae* | CFBP6466; A5423 | Martinique | *A. comosus* | Negative |
| *D. zeae* | PL47 | Hawaii, USA | *Brassica oleraceae* | Negative |
| *D. zeae* | PL65 | Hawaii, USA | *Colocasia esculenta* | Negative |
| *Enterobacter cloacae* | A5149 | Hawaii, USA | *Zingiber officinale* | Negative |
| *Enterobacter* sp. | PL247 | Hawaii, USA | Mizuna | Negative |
| *Enterobacter* sp. | PL168 | Hawaii, USA | *B. oleraceae* | Negative |
| *Erwinia amylovora* | A1084 | - | *Pyrus* spp. | Negative |
| *Erwinia* sp. | A5367 | Hawaii, USA | *Aglaonema* | Negative |
| *Klebsiella* sp. | A223 | - | *Vanda* spp. | Negative |
| *Musicola paradisiaca* | CFBP3696; A6064 | Cuba | *Musa* spp. | Negative |
| *Pantoea agglomerans* | A6222; DP138 | Wisconsin, USA | *Z. mays* | Negative |
| *P. agglomerans* | A6215 | Iowa, USA | *Z. mays* | Negative |
| *Pantoea* sp. | A1867 | Hawaii, USA | *C. papaya* | Negative |
| *Pectobacterium* sp. | PL34 | Hawaii, USA | *Hoodia* spp. | Negative |
| *Pectobacterium actinidiae* | LMG26003 | Jeju Korea, | *Actinidia chinensis* | Negative |
| *P. actinidiae* | LMG26004 | Jeju Korea, | *A. chinensis* | Negative |
| *P. aquaticum* | CFBP8637 | France | Environment/fresh water | Negative |
| *P. aroidearum* | LMG2414 | Israel | *Persea americana* | Negative |
| *P. atrosepticum* | LMG2374 | United Kingdom | *Apium graveolens* | Negative |
| *P. betavascularum* | LMG2466 | USA | *Beta vulgaris* | Negative |
| *P. betavascularum* | A6167 | California, USA | *B. vulgaris* | Negative |
| *P. brasiliense* | PL48 | Hawaii, USA | *B. oleraceae* | Negative |
| *P. brasiliense* | PL139 | Hawaii, USA | *B. oleraceae* | Negative |
| *P. brasiliense* | PL138 | Hawaii, USA | *B. oleraceae* | Negative |
| *P. brasiliense* | PL246 | Hawaii, USA | Pak choi | Negative |
| *P. brasiliense* | PL248 | Hawaii, USA | Mizuna | Negative |
| *P. brasiliense* | PL252 | Hawaii, USA | Radish | Negative |
| *P. brasiliense* | PL253 | Hawaii, USA | Radish | Negative |
| *P. cacticida* | LMG17936 | Arizona, United States | *Carnegiea gigantea* | Negative |
| *P. carotovorum* | LMG2404 | Denmark | *S. tuberosum* | Negative |
| *P. carotovorum* | A5280 | Hawaii, USA | Irrigation water | Negative |
| *P. carotovorum* | PL134 | Hawaii, USA | *B. oleraceae* | Negative |
| *P. carotovorum* | PL73 | Hawaii, USA | *S. tuberosum* | Negative |
| *P. fontis* | LMG30744 | Fresh water | Sungai waterfall Malaysia | Negative |
| *P. parmentieri* | LMG29774 | France | *S. tuberosum* | Negative |
| *P. parvum* | CFBP8631 | Finland | *S. tuberosum* | Negative |
| *P. polaris* | ICMP9180 | Netherlands | *S. tuberosum* | Negative |
| *P. polonicum* | LMG31077 | Gdansk, Poland | Ground water-potato fields | Negative |
| *P. punjabense* | LMG30622 | Punjab, Pakistan | *S. tuberosum* | Negative |
| *P. versatiles* | PL62 | Hawaii, USA | *S. tuberosum* | Negative |
| *P. wasabiae* | LMG8444 | Prefecture, Japan | *Eutrema wasabi* | Negative |
| *P. wasabiae* | A6159 | Wisconsin, USA | *S. tuberosum* | Negative |
| *P. zantedeschiae* | CFBP1357 | France | *Zantedeschia* spp. | Negative |
| *P. versatile* | ICMP9168 | Netherlands | *S. tuberosum* | Negative |
| *R. solanacearum* | A3450 | Trinidad | *S. lycopersicum* | Negative |
| *Rathayibacter rathayi* | LMG3717 | United Kingdom | *Dactylis glomerata* | Negative |
| *Stenotrophomonas* sp. | A5586 | Hawaii, USA | Anthurium | Negative |
| *Xanthomonas campestris*<br>pv. *leersiae* | NJ6.1.1 | Mali | *Leersia hexandra* | Negative |
| *X. citri* pv. *aracearum* | A6232 | Hawaii, USA | Syngonium | Negative |
| *X. phaseoli* pv.<br>*dieffenbachiae* | A5362 | Hawaii, USA | Anthurium | Negative |
| *X. sacchari* | A2111 | Hawaii, USA | *Colocasia esculenta* | Negative |
| *X. vesicatoria* | A3616 | South America | Tomato | Negative |
| **Exclusivity Panel (Plant/Soil samples)** |  |  |  |  |
| *Allium cepa* | Healthy onion | Hawaii, USA | - | Negative |
| *Colocasia esculenta* | Healthy taro | Hawaii, USA | - | Negative |
| *Phalaenopsis* sp. | Healthy orchid | Hawaii, USA | - | Negative |
| *Solanum tuberosum* | Healthy potato | Hawaii, USA | - | Negative |
| - | Non-infested<br>soil | Hawaii, USA | - | Negative |

Note: most of the DNA used in this study were taken from previous or ongoing studies in the laboratory (Dobhal et al, 2024; DeLude et al, 2022; Boluk et al, 2020).

### Limit of Detection (LOD)

The LOD was determined by two tests: a ten-fold serial dilution of pure genomic DNA from *D. dadantii* strain CFBP1269 and a ten-fold serial dilution of the same genomic DNA spiked with host crude lysate. Each LOD assessment was conducted independently by two operators, resulting in two trials for genomic DNA sensitivity and two trials for the assay combining bacterial genomic DNA with host crude lysate. In both the pure genomic DNA serial dilution and the spiked assays, *D. dadantii* was detectable down to 1 pg of genomic DNA. The presence of host DNA did not show any inhibitory effects on the LAMP reaction, as illustrated in Figure 3.

**Figure 3.**
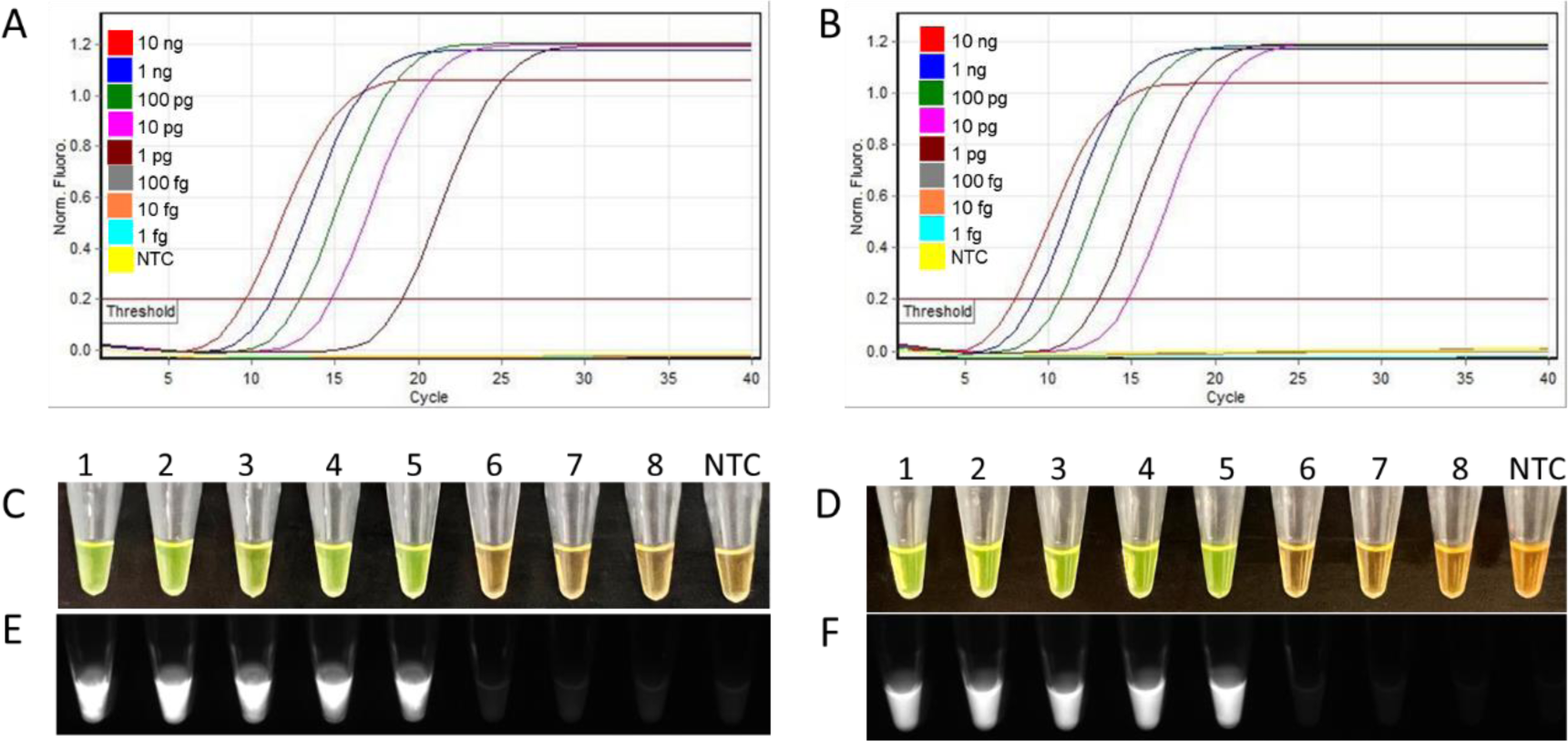
Limit of detection (LOD) for the *Dickeyadadantii*-specific LAMP assay determined using ten-fold serial dilutions of genomic DNA. (**A, C & E**) LOD assay with serially diluted *D. dadantii* genomic DNA in the following concentrations (left to right): 10 ng, 1 ng, 100 pg, 10 pg, 1 pg, 100 fg, 10 fg, 1 fg, and NTC (non-template control). (B, D, & F) LOD assay with serially diluted *D. dadantii* genomic DNA spiked with crude host lysate. A and B: sigmoidal amplification curves illustrating the LOD assay results. C and D: LAMP products after the addition of SYBR Green I dye, with amplification products turning bright green, and non-amplified samples remaining the original orange color. E and F: LAMP products with SYBR Green I dye under UV light; fluorescence indicates positive amplification.

### Multi-operator Validation Blind Test

To verify the repeatability of the developed diagnostic assay, multi-operator blind tests were conducted with eight samples. The results are represented in Figure 4. The results from all operators were consistent, further confirming the exclusivity of the developed diagnostic assay for detecting *D. dadantii* strains. No false positives or negatives were observed during these multi-operator blind tests.

**Figure 4.**
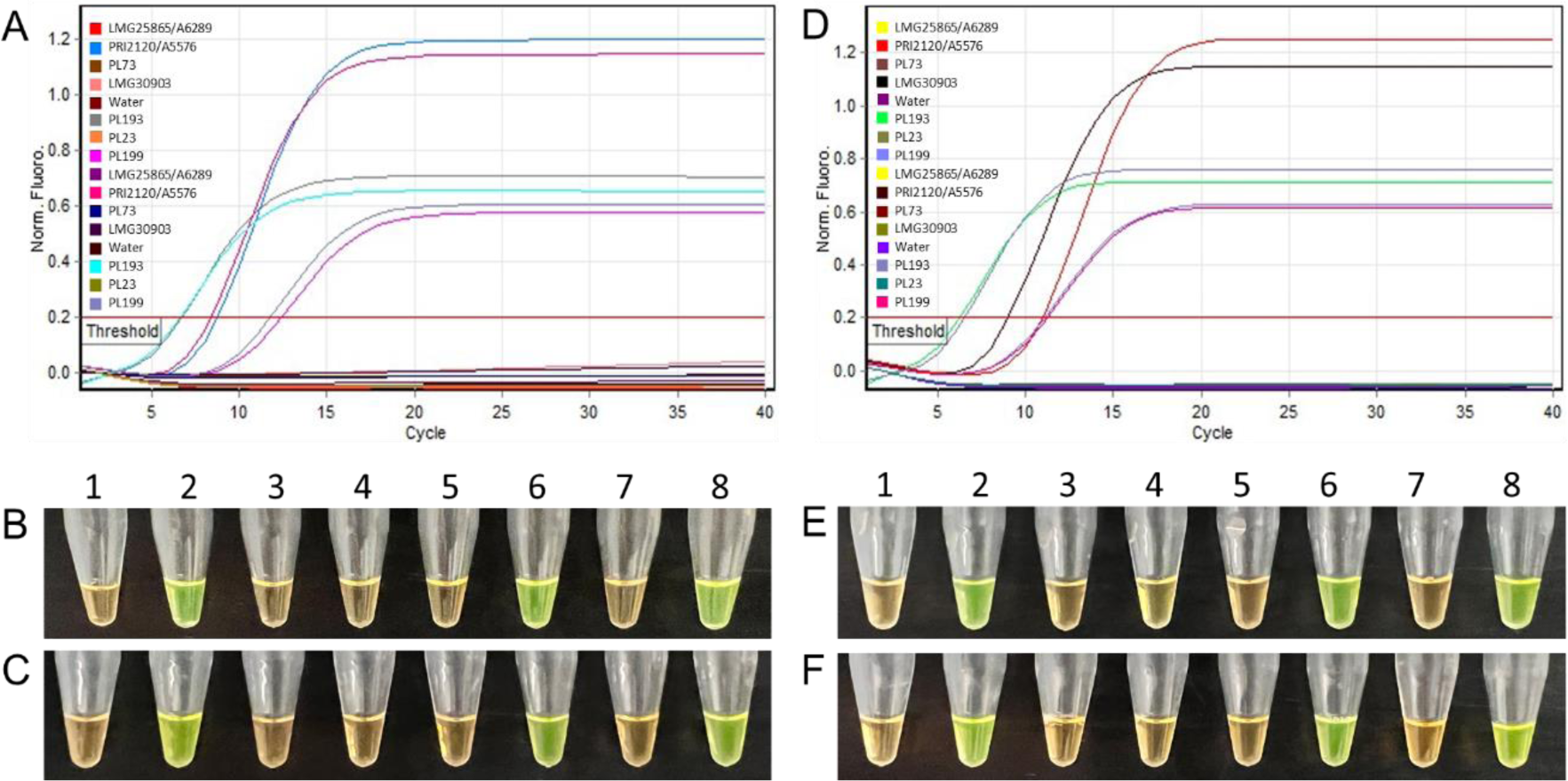
Multi-operator blind test assay results. (**A**) Sigmoidal amplification curves showing cycle threshold (CT) values from the LAMP assays performed by operators #1 and #2. (**B and C**) Visualization of the LAMP products for operators #1 and #2, respectively, after the addition of SYBR Green I dye; green color indicating amplification and orange indicating samples without amplification. (**D**) Quantification curve showing CT values from LAMP assays conducted by operators #3 and #4. (**E and F**) Visualization of LAMP products for operators #3 and #4, respectively, following the addition of SYBR Green I dye. All samples that tested positive were confirmed to contain *D. dadantii* genomic DNA. Tubes 1-8: *Dickeya solani* LMG25865/A6289*; D. dadantii* PRI2120/A5576; *Pectobacterium carotovorum* PL73; *D. undicola* LMG30903; Water; *D. dadantii* PL193; *D. dianthicola* PL23; *D. dadantii* PL199.

### Multi-instrument Detection

Multi-instrument detection and validation were carried out using three different instruments: Rotor-Gene Q, Bio-Rad thermocycler, and a dry bath platform. Eleven samples were tested with each instrument, including five strains of *D. dadantii* from the inclusivity panel, four strains of other species from the exclusivity panel, a DNA sample from healthy potato, and a non-template control (NTC) (Figure 5). The test results were visualized through amplification curves, melting curves, and the addition of SYBR Green I dye. All detection results were consistent, accurately reflecting the pathogenic association of the samples with no false negatives or false positives. The presence of single peak melting curves in positive reactions confirmed the specificity of the LAMP products. These outcomes demonstrate that the developed assay is robust and can be used effectively with any of the three instruments tested, whether in laboratory or in the field settings.

**Figure 5.**
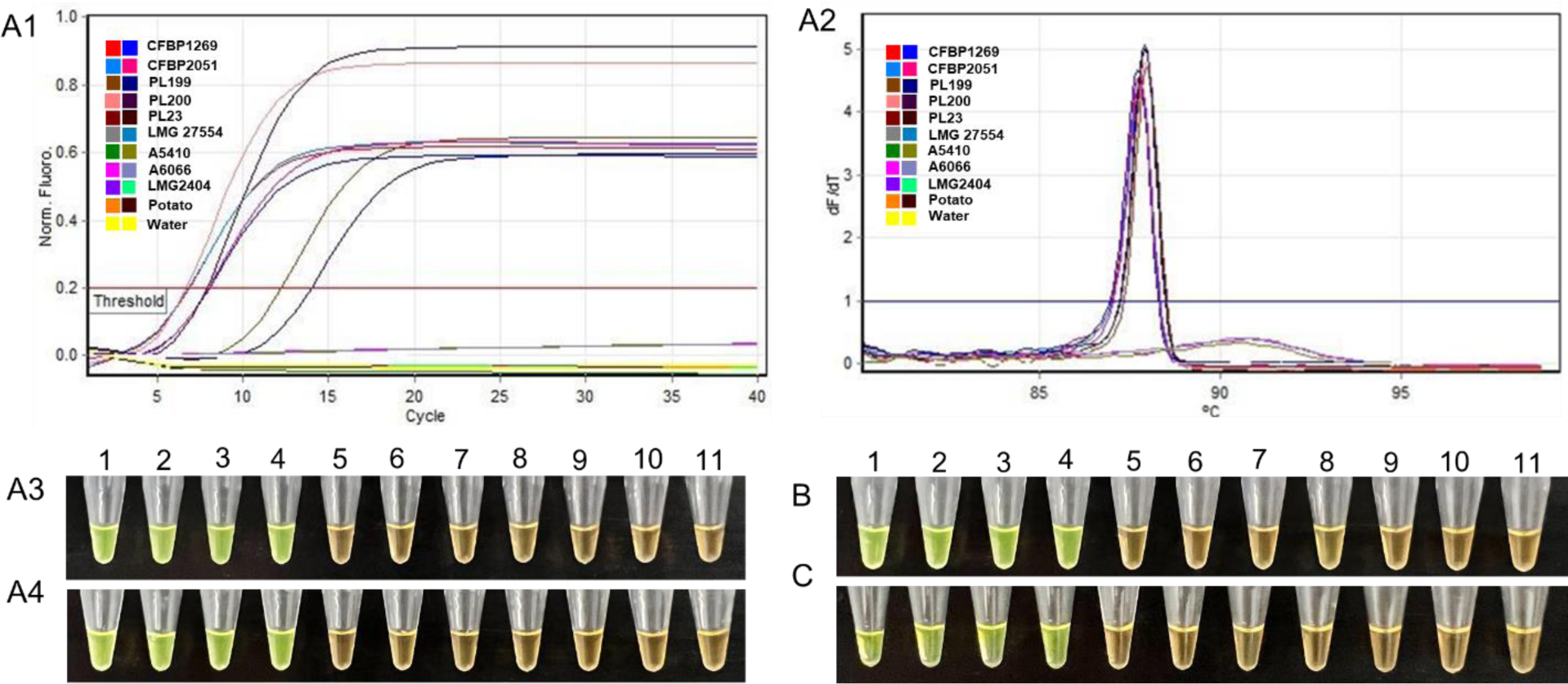
Validation of multi-device detection of *Dickeya dadantii* colonies using three different incubation platforms. Ten representative samples were tested. Tubes 1-11: *D. dadantii* strains CFBP1269, CFBP2051, PL199, PL200, *D. dianthicola* PL23, *D. solani* LMG 27554, *D. oryzae* A5410, *D. zeae* A6066, *Pectobacterium carotovorum* LMG2404, healthy potato crude lysate and non-template negative control (NTC; water), respectively. The results were plotted using sigmoidal amplification curves, melting curves, and SYBR Green I dye-based detection (A1-A4). Rotor-Gene Q amplification was performed independently by two groups, with positive results shown by sigmoidal curves and further verified by the addition of SYBR Green I dye; bright green color indicates positive amplification. (B) LAMP amplification using a Bio-Rad thermocycler and visualization with SYBR Green I dye; (C) LAMP amplification using a dry bath and visualization with SYBR Green I dye.

### Validation Using Artificially Infected Samples

To evaluate the diagnostic accuracy of the developed LAMP assay, potato slices inoculated with various *Dickeya* and *Pectobacterium* species, including *D. dadantii, D. dianthicola, D. zeae, D. solani, D. undicola,* and *P. punjabense,* were tested following the methodology described previously. The LAMP assay accurately identified the presence of the target pathogen, *D. dadantii*, in six infected samples with no cross-reaction in samples inoculated with other closely related species. No amplification was detected in the healthy potato lysate or the non-template control. The addition of SYBR Green I dye changed the color of the positively amplified samples from orange to green, confirming a positive result. Conversely, no color change was observed in the healthy potato or non-template water control (Figure 6).

**Figure 6.**
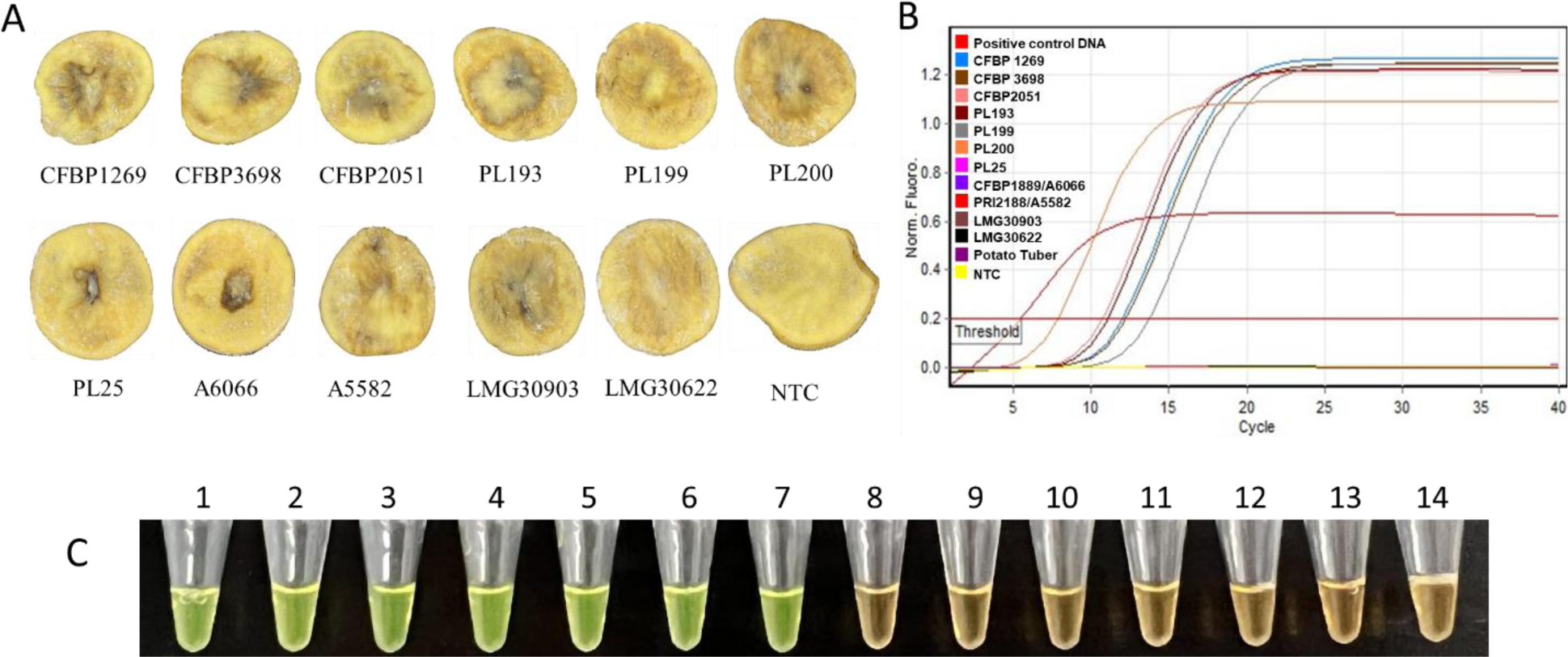
Detection of *Dickeya dadantii* from infected potato samples. (A) Infected potato slices infected with different *D. dadantii* isolates (CFBP1269, CFBP3698, CFBP2051, PL193, PL199, PL200), *D. dianthicola* (PL25), *D. zeae* (A6066)*, D. solani* (A5582)*, D. undicola* (LMG30903, *Pectobacterium punjabense* (LMG30622), and NC (negative) healthy potato. (B) Standard curve diagram– one *D. dadantii* positive control and six *D. dadantii-*infected potato slices showed amplification curves crossing Ct within 15 cycles (less than 10 minutes). No curve was observed for *D. dianthicola* (PL25), *D. zeae* (A6066)*, D. solani* (A5582)*, D. undicola* (LMG30903), *Pectobacterium punjabense* (LMG30622), NC (negative) healthy potato, or NTC (water). (C) Visualization of LAMP products after adding SYBR Green I dye; green color represents positive amplification (1-7) and orange color indicates negative results (8-14).

## DISCUSSION

An accurate and efficient diagnostic approach is crucial for effective disease management and for monitoring disease outbreaks. Such diagnostics play a pivotal role in early detection, enabling timely intervention and management, which can significantly reduce the spread of pathogen and disease, and mitigate their impact on plant health. In this study, we developed and optimized a LAMP assay for detecting *D. dadantii*, suitable for both field and laboratory settings. LAMP offers time and cost savings by enabling the production of up to 10^9^ copies of amplified DNA in under an hour (Soroka et al., 2021; Tomita et al., 2008). The method’s exceptional specificity is attributed to the use of two to three sets of primers (internal, external, and loop) that can recognize up to eight distinct sites on DNA or RNA targets (Soroka et al., 2021).

A robust assay necessitates accuracy and unfailing reliability for the success of a surveillance program, and its foundation rests on the precision of target selection (Arif et al, 2021; Dobhal et al., 2019, 2024). The availability of whole genomic data has enhanced the use of comparative genomic approaches to identify unique regions present in target species (Helmann et al., 2022; Domingo et al., 2021; Dobhal et al., 2020). The primers designed for the *D. dadantii*-specific LAMP assay targeted a unique region—the TetR/AcrR family transcriptional regulator coding sequence (CDS) and its flanking sequences—identified through comparative genomic analyses. This selected region is highly conserved and specific to *D. dadantii*, ensuring the longevity and robustness of the developed assay (Arif et al., 2021).

The *in silico* validation of primers is essential for the initial elimination of non-specific targets but does not guarantee the target’s *in vitro* specificity (Arif et al, 2021). Therefore, to ensure specificity, extensive inclusivity and exclusivity panels were developed. These included multiple strains of *D. dadantii* for inclusivity and various closely related species for exclusivity, as detailed in Table 2. Assays demonstrated that primers are reliable in detecting a broad range of *D. dadantii* strains and are unlikely to produce false negatives (Table 2; Figure 2). All bacterial strains and plant specimens included in the exclusivity panel yielded negative results. The exclusivity panel comprised closely related species and genera, common hosts, and bacteria that occupy similar ecological niches. These *in vitro* results demonstrate that the target sequence is highly specific to *D. dadantii*, confirming the assay’s capability to effectively distinguish between phylogenetically closely related species and other bacteria sharing the same niche or environment.

The accuracy of the assay hinges on its ability to detect low numbers of copies of the target region (Arif et al., 2021; Dobhal et al., 2020; Boluk et al., 2020). Failure to detect low copy numbers may result in false negatives, potentially allowing the undetected incursion of the target pathogen, which could lead to significant economic losses. The limit of detection for the developed LAMP assay was established through two independent sensitivity tests. The first test involved a 10-fold serial dilution of pure genomic *D. dadantii* DNA, while the second test used a 10-fold serial dilution of pure genomic *D. dadantii* DNA spiked with crude potato lysate. The assay consistently recognized *D. dadantii* at levels as low as 1 pg, equivalent to approximately 188 genome copies. These limits are comparable to those of other robust and sensitive assays, such as LAMP assay reported by Ocenar et al. (2019) for *D. dianthicola* and Domingo et al. (2021) for *P. parmentieri*. The spiked sensitivity test confirmed that there were no inhibitory effects from the host crude lysate on the assay’s performance, demonstrating its ability to maintain the same detection limits with or without the presence of host DNA (Dobhal et al, 2024).

The effectiveness of a diagnostic assay depends on its robustness and speed (Domingo et al., 2021; DeLude et al., 2022). The developed LAMP assay detected the target pathogen in less than 10 minutes, requiring minimal DNA preparation. This efficiency is attributed to its reduced susceptibility to plant inhibitors compared to PCR, enhancing its usability (Arif et al., 2021). The LAMP assay was validated by four independent operators (Figure 4) and was performed on multiple incubation platforms, including the Rotor-Gene Q, Bio-Rad thermocycler, and heating blocks (Figure 5). The obtained results were consistent, with no false positives or negatives reported. While tested with known infected samples, the assay detected all *D. dadantii* infected samples (Figure 6). The LAMP assay’s accessibility, accuracy, and robustness, along with its independence from complex laboratory equipment, significantly enhances its potential for pathogen detection and diagnosis. This flexibility renders it suitable for both laboratory research and field detection, offering considerable savings in economic and labor costs (Wong et al., 2018).

In conclusion, our research has yielded a sensitive, rapid, and easily field-deployable diagnostic LAMP assay for the specific detection of *D. dadantii*. This tool offers vast applications in detecting pathogen, managing diseases, conducting surveillance, and facilitating epidemiological studies, improving our ability to respond to agricultural challenges and safeguard crop health.

## MATERIALS AND METHODS

Plants and pathogen materials were used following national, international, and institutional guidelines.

### Target Genomic Region Selection

A genome region specific to *D. dadantii* was identified using the Progressive Mauve plugin Geneious Prime version 2022.1.1. The analysis included five genomes from different *D. dadantii* strains, 11 published genomes of other *Dickeya* species, and one from a *Pectobacterium* species, all retrieved from the National Center for Biotechnology Information (NCBI) genome database (see Data Availability). Approximately 800 local collinear blocks (LCBs) were identified among these genomes and were analyzed individually using Geneious Prime to pinpoint regions unique to *D. dadantii*. The sequence of the identified unique region was then subjected to a BLAST search on the NCBI database (https://blast.ncbi.nlm.nih.gov) to verify its exclusivity to *D. dadantii*. A nucleotide comparison ring image, illustrating the location of the region and its uniqueness across various *D. dadantii* strains, was created using the BLAST ring image generator (BRIG) (Alikhan et al., 2011). This genome comparison utilized the NCBI-BLAST version 2.13.0+ database, with *Dickeya dadantii* DSM1820 (NZ_CP023467) serving as the reference genome for the alignment.

### LAMP Primer Design

Six LAMP primers were designed targeting the unique gene TetR/AcrR family transcriptional regulator CDS and its adjacent sequence region. The primers, including the forward inner primer (DdntC-FIP), forward outer primer (DdntC-F3), backward inner primer (DdntC-BIP), backward outer primer (DdntC-B3), forward loop primer (Pp-LF), and backward loop primer (DdntC-LB), were designed using PrimerExplorer V5 (https://primerexplorer.jp/e/). The *in silico* specificity of each primer was verified against the available genome database using the NCBI GenBank BLASTn tool. Primers were synthesized by Genewiz (Azenta Life Sciences, South Plainfield, NJ) and are detailed in Table 1.

### Strains and DNA Extraction

Most of the strains used in this study were obtained from Pacific Culture Collection, Honolulu, HI. Glycerol stock cultures of bacterial strains stored at −80°C were streaked onto dextrose peptone agar (DPA, peptone 10 g 1^-1^, dextrose 5 g 1^-1^, and agar 17 g 1^-1^) or nutrient agar (NA) and incubated at 26-28°C for 1-2 days (Dobhal et al. 2020; Larrea-Sarmiento et al. 2018). Genomic DNA (gDNA) from each strain, as listed in Table 2, was either extracted using the Qiagen DNeasy Blood & Tissue Kit, following the manufacturer’s instructions (Qiagen, Germantown, MD) or taken from previous and other ongoing studies in the lab. Pure plant gDNA was extracted from 100 mg of healthy potato (*Solanum tuberosum*) tissue using the Qiagen DNeasy Plant Mini Kit (Qiagen). Crude lysate from plant material was extracted using the Plant Material Lysis Kit (Optigene, Sussex, UK) (Domingo et al, 2021).

### Specificity of LAMP Assay

The specificity of the LAMP assay was assessed using diverse strains from inclusivity and exclusivity panels (Table 2). Ninety non-target bacterial strains from various related species and genera, along with four different plant species (potato, taro, onion, and orchid), were included in the exclusivity panel. The inclusivity panel comprised nine different strains of *D. dadantii*: CFBP3698, PL193, PL199, PL200, A5419, A5643, A5576, and A5642from various regions, hosts, and collection years. *D. dadantii* strain CFBP1269 served as a positive control for all experiments, in accordance with protocols previously outlined by our lab (Domingo et al., 2021).

Each LAMP reaction had a total volume of 25 μL, consisting of 15 μL of Isothermal Master Mix (Optigene), 2 μL of the LAMP primer mix (1.6 μM each of DdntC-FIP and DdntC-B3, 0.2 μM each of DdntC-F3 and DdntC-B3, and 0.4 μM each of DdntC-LF and DdntC-LB), 1 μL of template DNA, and 7 μL of nuclease-free sterile water. The assays were conducted using a Rotor-Gene Q real-time thermocycler (Qiagen), with the *D. dadantii* type strain CFBP1269T as the positive control and a non-template (sterile water) reaction as the negative control. The amplification temperature was set at 65°C for 20 minutes, followed by a melting analysis from 80-99°C. Melting curves analysis was performed using Rotor-Gene Q software version 2.3.1.49 to identify any contamination or non-specific amplification. A volume of 3 μL of SYBR Green I (Life Technologies Corporation, Eugene, OR) was added to each tube after completion of the reaction, to observe fluorescence. The chemistry of SYBR Green I molecules enabled us to distinguish positive LAMP results, which appeared green under natural light and fluoresced under UV light, from negative results that remained orange under natural light without fluorescence under UV exposure.

### Determination of Limit of Detection (LOD)

Two tests were conducted to determine LOD, or sensitivity, of the developed LAMP assay for the specific detection of *D. dadantii*. Genomic DNA concentrations were quantified using the Qubit 4 fluorometer (Thermo Fisher Scientific, Waltham, MA). A ten-fold serial dilutions of pure gDNA from *D. dadantii* strain CFBP1269 in nuclease-free water were prepared, with final gDNA concentrations ranging from 1 nanogram (ng) to 1 femtogram (fg). A LAMP assay was then conducted following the previously described protocol. To assess inhibitor resistance and the impact of plant lysates on the assay, 5 μL crude lysate from healthy potato tuber tissue, extracted using the Plant Material Lysis Kit, was spiked into the serially diluted gDNA (1 ng to 1 fg) of *D. dadantii*. A negative control containing no bacterial gDNA, but only crude host plant lysate, was also included to assess primer specificity. Additionally, a non-template control (NTC) using nuclease-free water was included in each run to ensure no false-positive results were obtained.

### Multi-operator Validation Blind Test

Four independent operators carried out blind assays on a total of eight samples. These included three *D. dadantii* strains (PRI2120/A5576, PL193, PL199), one *D. dianthicola* strain (PL23), one *D. undicola* strain (LMG30903), one *D. solani* strain (LMG25865/A6289), one *P. carotovorum* strain (PL73), and one non-template control (NTC). Each operator followed the previously outlined LAMP protocol, and the results were subsequently compared with the initial diagnostic findings.

### Multi-instrument Validation

Multi-instrument detection was performed to assess the repeatability of the developed LAMP assay across three different incubation platforms: Rotor-Gene Q, Bio-Rad thermocycler, and dry bath, in accordance with the previously outlined LAMP assay instructions. The reaction conditions were standardized at 65℃ for 20 minutes for both the thermocyclers and dry bath. The test included a set of 11 samples comprising five strains of *D. dadantii* (CFPB1269, CFPB2051, PL199, PL200, PL23), three additional *Dickeya* species (*D. solani* LMG27554, *D. oryzae* A5410, *D. zeae* A6066), the type strain of *P. carotovorum* (LMG2404), healthy potato tuber gDNA, and a non-template control (NTC; water). These were amplified using each device. DNA samples were prepared following the extraction methods described earlier. Each LAMP reaction used 1 μl of the purified gDNA as a template. Results were assessed through amplification curves, melting curves, or SYBR Green dye.

### Validation with Infected Plant Tissues

Artificially infected potato samples were tested to validate the field applicability of the developed LAMP assay. The inoculum was prepared from overnight cultures. A total of six *D. dadantii* strains (CFBP1269 /A5416, CFBP3698/A6060, CFBP2051, PL193, PL199, PL200) and five strains from other species included in the exclusivity panel (*D. dianthicola* PL25, *D. zeae* 6066, *D. solani* A5582, *D. undicola* LMG30903, and *P. punjabense* LMG30622) were inoculated onto potato slices using the following method. Potato tubers were cleaned under tap water and immersed in a 0.6% sodium hypochlorite solution for 3 minutes. Afterward, the tubers were rinsed three times with sterile water and sliced. Approximately 10 μl of overnight-grown bacterial culture was inoculated into the flesh of each potato slice. The inoculated potato slices were placed into Petri dishes and incubated at room temperature for 12– 18 hours. Crude lysate was extracted from 100 mg of macerated tissue using the Plant Material Lysis Kit. Five μl of crude lysate was used in each LAMP reaction according to the protocol mentioned above. Genomic DNA of *D. dadantii*, healthy potato tuber lysate, and a nuclease-free water sample were used as the positive, negative, and non-template controls, respectively.

## Data Availability

The genomes were retrieved from the NCBI GenBank database and are available under the following accession numbers:

*Dickeya dadantii* 3937 (GCF_000147055.1)

https://www.ncbi.nlm.nih.gov/datasets/genome/GCF_000147055.1/

*Dickeya dadantii* DSM18020 (GCF_003049785.1)

https://www.ncbi.nlm.nih.gov/datasets/genome/GCF_003049785.1/

*Dickeya dadantii* FZ06 (GCF_025643375.1)

https://www.ncbi.nlm.nih.gov/datasets/genome/GCF_025643375.1/

*Dickeya dadantii* S3-1 (GCF_018904205.1)

https://www.ncbi.nlm.nih.gov/datasets/genome/GCF_018904205.1/

*Dickeya dadantii* M2-3 (GCF_020047155.1)

https://www.ncbi.nlm.nih.gov/datasets/genome/GCF_020047155.1/

*Dickeya solani* PPO 9019 (GCF_002846995.1)

https://www.ncbi.nlm.nih.gov/datasets/genome/GCF_002846995.1/

*Dickeya oryzae* ZYY5 (GCF_009372235.1)

https://www.ncbi.nlm.nih.gov/datasets/genome/GCF_009372235.1/

*Dickeya aquatica* 174/2 (GCF_900095885.1)

https://www.ncbi.nlm.nih.gov/datasets/genome/GCF_900095885.1/

*Dickeya fangzhongdai* DSM 101947 (GCF_002812485.1)

https://www.ncbi.nlm.nih.gov/datasets/genome/GCF_002812485.1/

*Dickeya zeae* MS2 (GCF_002887555.1)

https://www.ncbi.nlm.nih.gov/datasets/genome/GCF_002887555.1/

*Dickeya chrysanthemi* Ech1591 (GCF_000023565.1)

https://www.ncbi.nlm.nih.gov/datasets/genome/GCF_000023565.1/

*Dickeya lacustris* S29 (GCF_003934295.1)

https://www.ncbi.nlm.nih.gov/datasets/genome/GCF_003934295.1/

*Dickeya undicola* 2B12 (GCF_000784735.1)

https://www.ncbi.nlm.nih.gov/datasets/genome/GCF_000784735.1/

*Dickeya dianthicola* ME23 (GCA_003403135.1)

https://www.ncbi.nlm.nih.gov/datasets/genome/GCF_003403135.1/

*Dickeya dianthicola* 16MB01 (GCF_018361125.1)

https://www.ncbi.nlm.nih.gov/datasets/genome/GCF_018361125.1/

*Pectobacterium parmentieri* IFB5427 (GCF_003992745.1)

https://www.ncbi.nlm.nih.gov/datasets/genome/GCF_003992745.1/

## ACKNOWLEDGEMENTS

This work was funded by the USDA-ARS Agreement no. 58-2040-9-011, Systems Approaches to Improve Production and Quality of Specialty Crops Grown in the U.S. Pacific Basin; sub-project: Genome Informed Next Generation Detection Protocols for Pests and Pathogens of Specialty Crops in Hawaii. The Barry and Barbara Brennan Endowment, University of Hawaii, also supported this research. The bacterial strains utilized in this study were preserved and maintained through the National Science Foundation project (NSF-CSBR grant no. DBI-1561663). This research is a product of the course “PEPS/MBBE 627 Molecular Diagnostics: Principles and Practices.” The mention of specific trade names or commercial products in this publication does not constitute an official endorsement or recommendation by the University of Hawaii or the USDA.

## REFERENCES

1. Alikhan, N., Petty, N., Zakour, N., & Beatson, S. BLAST Ring Image Generator (BRIG): Simple prokaryote genome comparisons. BMC Genome, 1–10 (2011).

2. Arif, M., Busot, G.Y., Mann, R., Rodoni, B., and Stack, J.P. Field-deployable recombinase polymerase amplification assay for specific, sensitive and rapid detection of the US select agent and toxigenic bacterium, *Rathayibacter toxicus*. Biology 10: 620. doi10.3390/biology10070620 (2021).

3. Arif, M., Czajkowski, R., and Chapman, T. Editorial: Genome-wide analyses of *Pectobacterium* and *Dickeya* species. Front. Plant Sci. doi:10.3389/fpls.2022.855262 (2022).

4. Ash, G., Lang, J., Triplett, L., Stodart, B., Verdier, V., Cruz, C., Rott, P., Leach, J. Development of a genomics-Based LAMP (Loop-Mediated Isothermal Amplification) assay for detection of *Pseudomonas fuscovaginae* from rice. Plant disease, 909–915. 10.1094/PDIS-09-13-0957-RE (2014).

5. Boluk, G., and Arif, M. First report of *Dickeya dianthicola* as a causal agent of bacterial soft rot of potato in Hawaii. Plant Dis, doi10.1094/PDIS-11-18-2094-PDN (2019).

6. Boluk, G., Arizala, D., Dobhal, S., Zhang, J., Hu, J., Alvarez, A.M., and Arif, M. Genomic and phenotypic biology of novel strains of *Dickeya zeae* isolated from pineapple and taro in Hawaii: insights into genome plasticity, pathogenicity, and virulence determinants. Front Plant Sci. 12: 663851.doi: 10.3389/fpls.2021.663851 (2021).

7. Boluk, G., Dobhal, S., Crockford, A.B., Melzer, M. J., Alvarez, A. M. & Arif, M. Genome-informed recombinase polymerase amplification assay coupled with a lateral flow device for in-field detection of *Dickeya* species. Plant Dis. 10.1094/PDIS-09-19-1988-RE (2020).

8. Campos, H., & Ortiz, O. The Potato Crop: Its agricultural, nutritional, and social contribution to humankind. Switzerland: Springer. doi:10.1007/978-3-030-28683-5 (2020).

9. Cardoza, YF., Duarte V., and Lopes, C.A. First report of blackleg of potato caused by *Dickeya solani* in Brazil. Plant Dis. IOI, 243. doi: I0.1094/PDIS-07-16-1045-PDN (2016).

10. CIP. Potato facts and Figures. Retrieved from International Potato Center: https://cipotato.org/potato (2022).

11. Charkowski, A., et al. The role of secretion systems and small molecules in soft-rot *Enterobacteriaceae* pathogenicity. Annu Rev Phytopathol 50: 425–449 (2012).

12. Condemine, G., & Le Derout, B. Identification of new *Dickeya dadantii* virulence factors secreted by the type 2 secretion system. Plos One (2022).

13. DeLude, A. et al. Loop-mediated isothermal amplification (LAMP) assay for specific and rapid detection of *Dickeya fangzhongdai* targeting a unique genomic region. Scientific reports (12). 10.1038/s41598-022-22023-4 (2022).

14. Dobhal, S. et al. Comparative genomics reveals signature regions used to develop a robust and sensitive multiplex TaqMan real-time qPCR assay to detect the genus *Dickeya* and *Dickeya dianthicola*. Journal of applied microbiology, 1703-1719. 10.1111/jam.14579 (2020).

15. Dobhal, S., Santillana, G., Stulberg, M.J., Arizala, D., Alvarez, A.M., and Arif, M. Development and validation of genome-informed and multigene-based qPCR and LAMP assays for accurate detection of Dickeya solani: a critical quarantine pathogen threatening potato industry. doi: 10.1101/2024.03.21.586178 (2024).

16. Domingo, R. et al. Genome-informed loop-mediated isothermal amplification assay for specific detection of *Pectobacterium parmentieri* in infected potato tissues and soil. Scientific reports. doi: 10.1038/s41598-021-01196-4 (2021).

17. Glasner, J. et al. Genome sequence of the plant-pathogenic bacterium *Dickeya dadantii* 3937. J. Bacteriol. 10.1128/jb.01513-10 (2011).

18. He, W. et al. Three highly sensitive and high-throughput serological approaches for detecting *Dickeya dadantii* in sweet potato. Plant disease. 105(4), 832–839. 10.1094/PDIS-07-20-1551-RE (2021).

19. Helmann, T., Filiatrault, M., & Stodghill, P. Genome-wide identification of genes important for growth of *Dickeya dadantii* and *Dickeya dianthiocola* in potato (*Solanum tuberosum*) Tubers. Front Microbiology, 13 (2022).

20. HLPE. Impact of COVID-19 on food security and nutrition: Developing effective policy responses to address the hunger and malnutrition pandemic. FAO. Retrieved from https://www.fao.org/3/cb1000en/cb1000en.pdf (2020).

21. Hugouvieux-Cotte-Pattat, N. & Van Gijsegem, F. Diversity within the *Dickeya zeae* complex, identification of *Dickeya zeae* and *Dickeya oryzae* members, proposal of the novel species *Dickeya parazeae* sp. nov. Int J Syst Evol Microbiol. 71, 5059 (2021).

22. Hugouvieux-Cotte-Pattat, N. Metabolism and virulence strategies in *Dickeya*–host interactions. Progress in molecular biology and translational science, 93–129 (2016).

23. Ivanov, A., Safenkova, I., Zherdev, A., & Dzantiev, B. The potential use of isothermal amplification assays for in-field diagnostics of plant pathogens. Plants. 10.3390/plants10112424 (2021).

24. Larrea-Sarmiento, A., Dhakal, U., Boluk, G., Fatdal, L., Alvarez, A., Strayer-Scherer, A., & Arif, M. Development of a genome-informed loop-mediated isothermal amplification assay for rapid and specific detection of *Xanthomonas euvesicatoria*. Scientific Reports. 10.1038/s41598-018-32295-4 (2018).

25. Li, M. et al. Development and clinical application of a rapid and visual loop-mediated isothermal amplification test for tetM gene in *Clostridioides difficile* strains cultured from feces. Int J Infect Dis, 676–684. doi: 10.1016/j.ijid.2022.07.032 (2022).

26. Liu, Q., Xiao, W., Wu, Z., Li, S., Yuan, Y., & Li, H. Identification of *Dickeya dadantii* as a causal agent of Banana bacterial sheath rot in China. Journal of Plant pathology, 98(3), 503–510. 10.1094/PDIS-07-13-0711-RE (2016).

27. Mansfield, J. et al. Top 10 plant pathogenic bacteria in molecular plant pathology. Molecular plant pathology, 13(6), 614–629. 10.1111/j.1364-3703.2012.00804.x (2012).

28. Mori, Y., & Notomi, T. Loop-mediated isothermal amplification (LAMP): a rapid, accurate, and cost-effective diagnostic method for infectious diseases. J. Infect. Chemother, 62–69 (2009).

29. Notomi, T., Okayama, H., Masubichi, H., Yonekawa, T., Watanabe, K., Amino, N., & Hase, T. Loop-mediated isothermal amplification of DNA. Nucleic acids research. 10.1093/nar/28.12.e63 (2000).

30. Ocenar, J. et al. Development of a robust, field-deployable loop-mediated isothermal amplification (LAMP) assay for specific detection of potato pathogen *Dickeya dianthicola* targeting a unique genomic region. Plos One. 10.1371/journal.pone.0218868 (2019).

31. Parida, M., Sannarangaiah, S., Dash, P., Rao, P., & Morita, K. Loop mediated isothermal amplification (LAMP): a new generation of innovative gene amplification technique; perspectives in clinical diagnosis of infectious diseases. Rev. Med. Virolo, 407–421 (2008).

32. Pritchard, L., Humphris, S., Saddler, G. S., Parkinson, N. M., Bertrand, V., Elphinstone, J. G. & Toth, I. K. Detection of phytopathogens of the genus *Dickeya* using a PCR primer prediction pipeline for draft bacterial genome sequences. Plant pathology 62(3), 587–596 (2013).

33. Safenkova, I. et al. Development of a lateral flow immunoassay for rapid diagnosis of potato blackleg caused by *Dickeya* species. Analytical and bioanalytical chemistry, 1915-1927. 10.1007/s00216-016-0140-6 (2017).

34. Samson, R., Legendre, J., Christen, R., Fischer-Le Saux, M., Achouak, W., & Gardan, L. Transfer of *Pectobacterium chrysanthemi* (Burkholder et al. 1953) Brenner et al. 1973 and *Brenneria paradisiaca* to the genus *Dickeya* gen. nov. as *Dickeya chrysanthemi* comb. nov. and *Dickeya paradisiaca* comb. nov. and delineation of four novel species, International journal of systematic and evolutionary microbiology (2005).

35. Savary, S., Ficke, A., Aubertot, J., & Hollier, C. Crop losses due to diseases and their implications for global food production losses and food security. Food Secur 4, 512–537 (2012).

36. Soleimani-Delfan, A., Etemadifar, Z., Emtiazi, G., & Bouzari, M. Isolation of *Dickeya dadantii* strains from potato disease and biocontrol by their bacteriophages. Braz. J. Microbiol, 46, 791–797 (2015).

37. Soroka, M., Wasowicz, B., & Rymaszewska, A. Loop-mediated isothermal amplification (LAMP): The better sibling of PCR? Cells (2021).

38. Tomita, N., Mori, Y., Kanda, H., & Notomi, T. Loop-mediated isothermal amplification (LAMP) of gene sequences and simple visual detection of products. Nat. Protoc, 877–882 (2008).

39. Wang, X., et al. *Dickeya oryzae* sp. nov., isolated from the roots of rice. Int J Syst Evol Microbiol. 70(7), 4171–4178 (2020).

40. Wong, Y., Othman, S., Lau, Y., Radu, S., & Chee, H. Loop-mediated isothermal amplification (LAMP): a versatile technique for detection of microorganisms. J. Appl. Microbiol., 626–643 (2018).

41. Youdkes, D., Helman, Y., Burdman, S., Matan, O., & Jurkevitch, E. Potential control of potato soft rot disease by the obligate predators *Bdellovibrio* and like organisms. Appl. Environ. Microbiol, 86. doi: 10.1128/AEM.02543-19 (2020).

42. Zou, Y., Mason, M., & Botella, J. Evaluation and improvement of isothermal amplification methods for point-of-need plant disease diagnostics. Plos One. (2020).

